# Rapid adaptation to Elevated Extracellular Potassium in the Pyloric Circuit of the Crab, *Cancer borealis*

**DOI:** 10.1101/636910

**Authors:** Lily S. He, Mara C.P. Rue, Ekaterina O. Morozova, Daniel J. Powell, Eric J. James, Manaswini Kar, Eve Marder

**Affiliations:** Biology Department and Volen Center, Brandeis University, Waltham, MA 02454; Program in Neuroscience, Harvard University, Boston, MA 02115; Biology Department, Adelphi University, Garden City, NY 11530; Department of Neurobiology, University of Pittsburgh, Pittsburgh, PA 15213

## Abstract

Elevated [K^+^] is often used to alter excitability in neurons and networks by shifting the potassium equilibrium potential (E_K_) and, consequently, the resting membrane potential. We studied the effects of increased extracellular [K^+^] on the well-described pyloric circuit of the crab, *Cancer borealis.* A 2.5-fold increase in extracellular [K^+^] (2.5x[K^+^]) depolarized Pyloric Dilator (PD) neurons and resulted in short-term loss of their normal bursting activity. This period of silence was followed within 5-10 minutes by the recovery of spiking and/or bursting activity during continued superfusion of 2.5x[K^+^] saline. In contrast, when PD neurons were pharmacologically isolated from pyloric presynaptic inputs, they exhibited no transient loss of spiking activity in 2.5x[K^+^], suggesting the presence of an acute inhibitory effect mediated by circuit interactions. Action potential threshold in PD neurons hyperpolarized during an hour-long exposure to 2.5x[K^+^] concurrent with the recovery of spiking and/or bursting activity. Thus, the initial loss of activity appears to be mediated by synaptic interactions within the network, but the secondary adaptation depends on changes in the intrinsic excitability of the pacemaker neurons. The complex sequence of events in the responses of pyloric neurons to elevated [K^+^] demonstrates that electrophysiological recordings are necessary to determine both the transient and longer-term effects of even modest alterations of K^+^ concentrations on neuronal activity.

**Significance Statement:** Solutions with elevated extracellular potassium are commonly used as a depolarizing stimulus. Moreover, hyperkalemia is associated with a number of disease states, including epileptic seizures and brain traumas. We studied the effects of high [K^+^] saline on the well-described pyloric circuit of the crab stomatogastric ganglion. A 2.5-fold increase in extracellular [K^+^] led to a transient loss of activity in pyloric neurons that was not due to depolarization block. This was followed by a rapid increase in excitability and concurrent recovery of spiking and rhythmic bursting activity within minutes. These results suggest that effects of high [K^+^] on neuronal circuits can be complex and non-stationary.

## Introduction

Neuronal circuits must be robust to various environmental challenges. This is especially true for central pattern generators (CPGs) that produce essential motor patterns such as breathing, walking, and chewing (Marder and Calabrese, 1996). Maintaining stability over a range of perturbations involves multiple intrinsic and synaptic mechanisms that operate over the course of minutes to days (Von Euler, 1983; Marder and Bucher, 2001; Harris-Warrick, 2010). Moreover, both theoretical and experimental evidence suggest that robust CPGs with similar activity patterns can have widely variable underlying cell intrinsic and synaptic conductances (Prinz et al., 2004; Marder and Goaillard, 2006; Schulz et al., 2006; Schulz et al., 2007; Goaillard et al., 2009; Norris et al., 2011; Roffman et al., 2011). Nevertheless, these individually variable circuits must maintain reliable outputs. How such circuits respond and adapt to environmental challenges remains an open question.

There are a number of global perturbations such as changes in temperature or pH that are likely to influence the behavior of most, if not all, neurons in a circuit. Previous work on the pyloric circuit of the crab stomatogastric ganglion (STG) has shown that the pyloric rhythm is remarkably resilient to substantial changes in temperature (Tang et al., 2010) or pH (Haley et al., 2018). Elevated extracellular potassium concentration ([K^+^]) is a physiologically relevant depolarizing stimulus associated with a wide array of conditions including thermal stress, epileptic seizures, kidney failure, traumatic brain injury, and stroke (Baylor and Nicholls, 1969; Katayama et al., 1990; Pérez-Pinzón et al., 1995; Jensen and Yaari, 1997; Rodgers et al., 2007; Krishnan and Kiernan, 2009; Morrison III et al., 2011; Arnold et al., 2014; Chauvette et al., 2016).

Experimentally, increased extracellular [K^+^] is often used to depolarize neurons and networks, with the usual aim of increasing neuronal and network activity (Ballerini et al., 1999; Lin et al., 2008; Panaitescu et al., 2009; Ruangkittisakul et al., 2011; Rybak et al., 2014; Sharma et al., 2015). In this study, we describe a series of short and longer-term effects of elevated [K^+^] on the pyloric circuit of the STG. In elevated [K^+^], we observe a short-term period (several minutes) in which rhythmic network activity is lost or slowed, followed by a period of adaptation in which rhythmic activity returns while still in elevated [K^+^]. These results demonstrate that the effects of high [K^+^] on neurons and networks can be complex and non-stationary.

## Materials and Methods

### Animals and dissections

Adult male Jonah Crabs, *Cancer borealis*, (N = 81) were obtained from Commercial Lobster (Boston, MA) between December 2016 and December 2019 and maintained in artificial seawater at 10-12°C in a 12-hour light/dark cycle. On average, animals were acclimated at this temperature for one week before use. Prior to dissection, animals were placed on ice for at least 30 minutes. Dissections were performed as previously described (Gutierrez and Grashow, 2009). In short, the stomach was dissected from the animal and the intact stomatogastric nervous system (STNS) was removed from the stomach including the commissural ganglia, esophageal ganglion and stomatogastric ganglion (STG) with connecting motor nerves. The STNS was pinned in a Sylgard-coated (Dow Corning) dish and continuously superfused with 11°C saline.

### Solutions

Physiological (control) *Cancer borealis* saline was composed of 440 mM NaCl, 11 mM KCl, 26 mM MgCl2, 13 mM CaCl2, 11 mM Trizma base, 5 mM maleic acid, pH 7.4-7.5 at 23°C (approximately 7.7-7.8 pH at 11°C). High - 1.5x, 2x, 2.5x, and 3x[K^+^] - salines (16.5, 22, 27.5 and 33mM KCl respectively) were prepared by adding more KCl salt to the normal saline. 10^-5^M Picrotoxin (PTX) was used to block inhibitory glutamatergic synapses (Marder and Eisen, 1984). 10^-7^M Tetrodotoxin (TTX) was used to block voltage-gated sodium channels for measurements of graded inhibition.

### Electrophysiology

Intracellular recordings from STG somata were made in the desheathed STG with 10–30 MΩ sharp glass microelectrodes filled with internal solution (10 mM MgCl_2_, 400 mM potassium gluconate, 10 mM HEPES buffer, 15 mM NaSO_4_, 20 mM NaCl (Hooper et al., 2015)). Intracellular signals were amplified with an Axoclamp 900A amplifier (Molecular Devices, San Jose). Extracellular nerve recordings were made by building wells around nerves using a mixture of Vaseline and mineral oil and placing stainless-steel pin electrodes within the wells to monitor spiking activity. Extracellular nerve recordings were amplified using model 3500 extracellular amplifiers (A-M Systems). Data were acquired using a Digidata 1440 digitizer (Molecular Devices, San Jose) and pClamp data acquisition software (Molecular Devices, San Jose, version 10.5). For identification of Pyloric Dilator (PD) and Lateral Pyloric (LP) neurons, somatic intracellular recordings were matched to action potentials on the pyloric dilator nerve (*pdn*), lateral pyloric nerve (*lpn*) and/or the lateral ventricular nerve (*lvn*).

### Elevated [K^+^] saline application

Prior to applications of elevated [K^+^] saline, baseline activity was recorded for 30 minutes in control saline. Following the baseline recording, the STNS was superfused with elevated [K^+^] saline in concentrations ranging from 1.5x to 3x [K^+^] for 90 minutes. The preparation was then superfused with control saline for 30 minutes. Recordings from PD neurons in the isolated pacemaker kernel were made by superfusing the preparation with 10^-5^M Picrotoxin (PTX) saline until the inhibitory synaptic potentials in the PD neurons disappeared (20 minutes). These preparations were then exposed to 2.5x[K^+^] PTX saline for 90 minutes, followed by a 30-minute wash in PTX saline.

### Measuring synaptic strength between LP and PD neurons

We measured the strength of graded synaptic inhibition from the LP neuron onto PD neurons and from the PD neuron onto LP neurons in physiological saline with 10^-7^ M TTX and over time in 2.5x[K^+^] TTX saline. The synaptic currents were measured using two different protocols: voltage clamp and current clamp.

Voltage clamp protocol: The LP neuron and a PD neuron were both held in two-electrode voltage clamp. The synaptic current from the LP neuron to the PD neuron was measured as the current elicited in the postsynaptic neuron, held at either -90mV or - 10mV, in response to depolarizing the presynaptic neuron to -10mV for 1-second steps. The synaptic current was measured at the peak of the postsynaptic response. The reversal potential of the synaptic current was estimated by fitting a line to the peak postsynaptic currents at -90mV and -10 mV. The synaptic current from the PD neuron to the LP neuron was measured as the current elicited in the postsynaptic neuron, held at -50 mV, in response to depolarizing the presynaptic neuron to -10mV. These measurements were repeated every 5 minutes for the entire experiment, which consisted of a 30-minute baseline in 10^-7^M TTX saline followed by 60 minutes in 2.5x[K^+^] TTX saline and finally a 30-minute wash in TTX saline.

Current clamp protocol: We used two-electrode current clamp to inject a series of ten 2-second current steps into the LP neuron from -50mV up to -20mV while simultaneously recording from a PD neuron held at -50mV. These current steps were repeated every 3 minutes for the entire experiment. We recorded the synaptic responses for 30 minutes in TTX saline. The preparation was then superfused with 2.5x[K+] TTX saline for 90 minutes, followed by a 30-minute wash in TTX saline.

### Threshold and excitability measurements

To measure the action potential threshold and excitability of PD neurons, two-electrode current clamp was used to apply slow ramps of current from -4nA to +2nA over 60 seconds. Resting membrane potential and input resistance were measured during baseline recordings and after the application of elevated [K^+^] to ensure the integrity of the preparation (neurons with input resistance <4MΩ were discarded). Three ramps were performed during baseline recordings at 10-minute intervals, and ramps were performed in 2.5x[K^+^] at 5, 10, 20, 30, 40, 50, 60, 70, 80 and 90 minutes after the start of elevated [K^+^] superfusion. After the preparation was returned to control saline, three ramps were performed again at 10-minute intervals. In recordings from the PD neurons with glutamatergic synapses blocked by PTX, baseline ramps were performed as described above, followed by three ramps in PTX saline. Preparations were then superfused with 2.5x[K^+^] PTX saline and washed in PTX saline following the same ramp procedure.

### Data acquisition and analysis

Recordings were acquired using Clampex software (pClamp Suite by Molecular Devices, San Jose, version 10.5) and visualized and analyzed using custom MATLAB analysis scripts. These scripts were used to detect and measure voltage response amplitudes and membrane potentials, plot raw recordings and processed data, generate spectrograms, and perform some statistical analyses.

### Spectral analysis

Spectrograms were calculated using the Burg (1967) method for estimation of the power spectrum density in each time-window. The Burg method (1967) fits the autoregressive (AR) model of a specified order p in the time series by minimizing the sum of squares of the residuals. The fast-Fourier transform (FFT) spectrum is estimated using the previously calculated AR coefficients. This method is characterized by higher resolution in the frequency domain than traditional FFT spectral analysis, especially for a relatively short time window (Buttkus, 2000). We used the following parameters for the spectral estimation: data window of 3.2 s, 50% overlap to calculate the spectrogram, number of estimated AR-coefficients p=(window/4)+1. Before the analysis, voltage traces were low pass filtered at 5 Hz using a six-order Butterworth filter and down-sampled. PD neuron burst frequency was calculated as the mean frequency at the peak spectral power in each sliding window.

### Analysis of interspike interval distributions

Intracellular voltage traces were thresholded to obtain spike times. Distributions of inter-spike intervals (ISIs) were calculated within 2-minute bins. Hartigan’s dip test of unimodality (Hartigan and Hartigan, 1985) was used to obtain the dip statistic for each of these distributions. This dip statistic was compared to Table 1 in Hartigan and Hartigan (1985) to find the probability of multi-modality. The test creates a unimodal distribution function that has the smallest value deviations from the experimental distribution function. The largest of these deviations is the dip statistic. The dip statistic shows the probability of the experimental distribution function being bimodal. Larger value dips indicate that the empirical data are more likely to have multiple modes (Hartigan and Hartigan, 1985).

### Activity pattern plots

For all recordings, we determined the time of spikes over the course of the experiment. For the recovery time plots, silence was defined as no more than 2 spikes in a 30-second sliding window. Otherwise, all spike behaviors (even if irregular) were counted as active spiking.

To determine more broadly the activity pattern of each PD neuron across the experiment, we analyzed the distribution of log-transformed ISIs in 2-minute bins using Hartigan’s dip statistic. If the dip statistic was 0.05 or higher the neuron was considered to be bursting. If the dip statistic was lower than 0.05 the neuron was considered to be tonically firing. Neurons with some spikes, but not enough ISIs to calculate the dip, were classified as sparsely firing. Neurons with no spikes in the observed window were classified as silent. We then plotted the activity patterns in these four categories – bursting, tonic, sparse firing and silent – for each PD neuron across the entire experiment.

### Identification of the spike threshold

The spike threshold was identified as the voltage point of the maximum curvature before the first spike. To find this, we calculated the first derivative of the voltage (dV/dt) and defined the spike onset as the point when dV/dt crosses the threshold value of 10 mV/ms.

All electrophysiology analysis scripts are available at the Marder lab GitHub (https://github.com/marderlab).

### Statistics

Statistical analysis was carried out using Excel 365 for ANOVA tests, and MATLAB 2018a for all other analyses as described above.

## Results

### Network activity in the pyloric circuit

The STNS of the crab *Cancer borealis* was isolated intact from the stomach and pinned in a dish, allowing us to change the composition of the continuously flowing superfused saline (Fig. 1*A*). The STG contains identified neurons that drive the pyloric rhythm which filters food through the animal’s foregut. Figure 1*B* shows a recording from the lateral ventricular nerve (*lvn)* which contains the axons of the LP, pyloric (PY) and PD neurons; the stereotypical triphasic pyloric pattern consists of repeated bursts of activity from the LP, PY and PD neurons. Figure 1*C* illustrates the connectivity diagram of the pyloric network. The anterior burster (AB) neuron, the intrinsic oscillator that drives the circuit, is strongly electrically coupled to the two PD neurons, which burst synchronously with the AB neuron. Together the AB and two PD neurons form the pacemaker kernel of the network, and their coordinated bursting initiates each triphasic cycle (Maynard, 1972). Glutamatergic and cholinergic synaptic connections between neurons in the STG are all inhibitory and are both graded and spike mediated (Graubard et al., 1980; Manor et al., 1997). LP and PY neurons’ bursting activity, which results from post-inhibitory rebound, compose the second and third phases of the pyloric rhythm respectively, due to rhythmic inhibition from the pacemaker and reciprocal inhibition between LP and PY. (Hartline and Gassie, 1979; Selverston and Miller, 1980).

**Figure 1.**
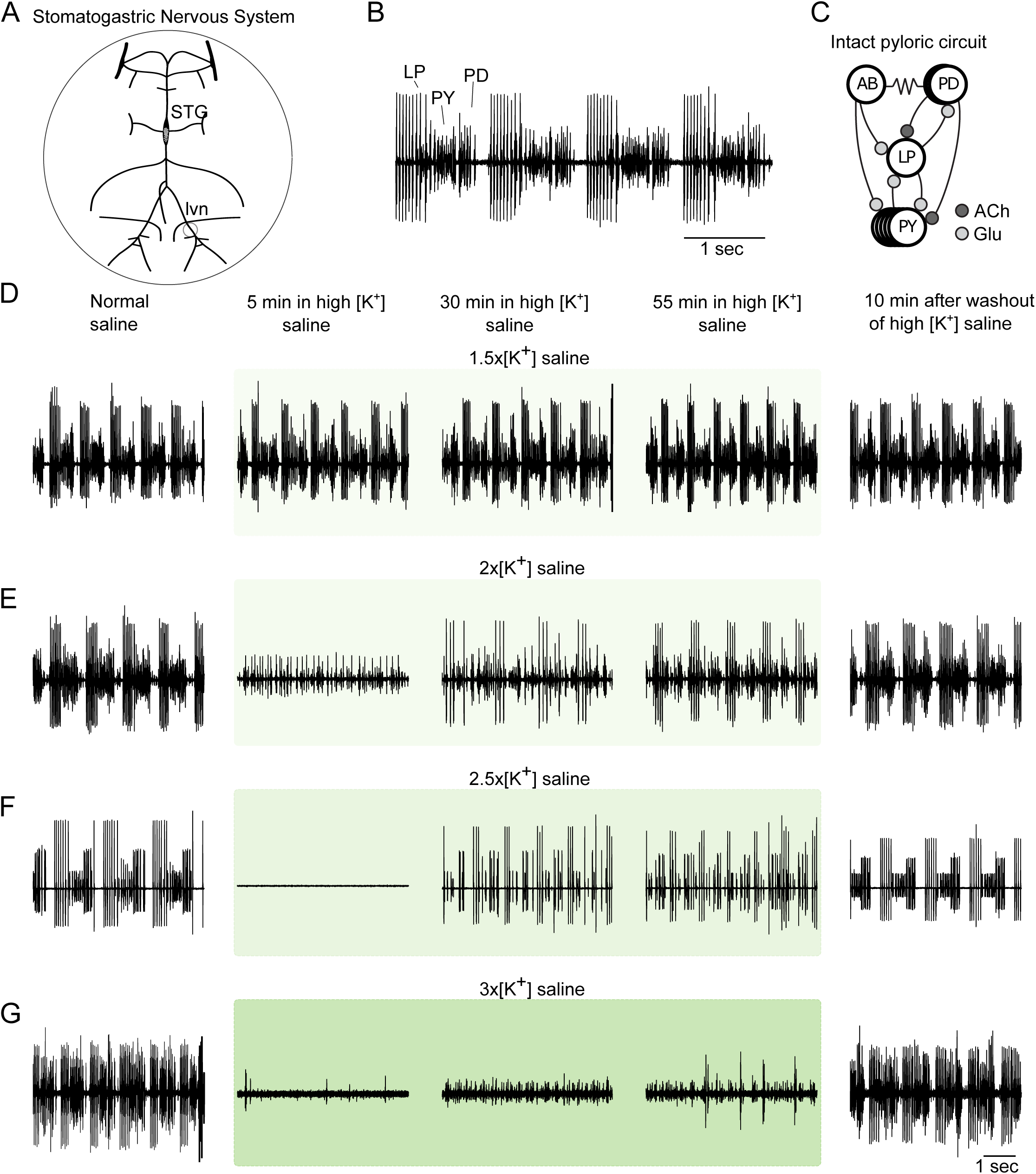
The pyloric rhythm is disrupted by large changes in extracellular [K^+^]. (A) Diagram of the dissected stomatogastric nervous system (STNS) showing the stomatogastric ganglion (STG) and motor nerves (*lvn*). The entire STNS was superfused with saline with altered [K^+^]. (B) The triphasic pyloric rhythm is illustrated in an extracellular recording from the *lvn,* which contains axons from LP, PY and PD neurons. (C) Connectivity diagram of the pyloric circuit of the crab *Cancer borealis*. All chemical synapses are inhibitory. Resistor symbols denote electrical synapses (D - G) Recordings of *lvn* spiking activity during application of elevated [K^+^] saline (green boxes) with concentrations 1.5x, 2.0x, 2,5x and 3.0x the physiological extracellular concentration of potassium.

### The pyloric rhythm is temporarily disrupted by high extracellular potassium

To test the response of the pyloric rhythm to changes in extracellular [K^+^], we superfused saline with elevated [K^+^] over the STNS while continuously recording the activity of pyloric neurons extracellularly from the *lvn*. We tested concentrations of [K^+^] that were 1.5, 2, 2.5 and 3-times the physiological concentrations to study the dose-dependent responses of pyloric neurons to changes in extracellular [K^+^]. When extracellular [K^+^] was increased to 1.5x the physiological concentration (Fig. 1*D*, N = 4), the pyloric rhythm remained triphasic and was negligibly affected. When exposed to 2x [K^+^] (Fig. 1*E, N* = 20), the response of pyloric neurons was more variable. In some cases (N = 9/20) there was a short disruption of the triphasic pyloric rhythm, which was followed by the recovery of spiking activity.

Higher concentrations of extracellular [K^+^] produced more pronounced effects on pyloric activity. 2.5x [K^+^] (Fig. 1*F, N* = 12 extracellular only recordings) reliably and profoundly altered the pyloric rhythm. During the application of 2.5x[K^+^] saline, all preparations exhibited a disruption of action potentials from pyloric neurons, followed by recovery of spiking activity during continued exposure to 2.5x[K^+^] saline. Application of 3x[K^+^] saline to the STNS resulted in consistent loss of the pyloric rhythm (Fig. 1*G, N* = 5). Nonetheless, at 3x[K^+^], few of the preparations recovered sustained activity. Based on these responses, we settled on 2.5x[K^+^] for further study, as it reliably disrupted the pyloric rhythm and was accompanied by a sustained recovery of spiking or bursting activity during the continued application of 2.5x[K^+^] saline.

The pattern of loss and recovery of pyloric activity in 2.5x[K^+^] saline was consistent across all experiments. To obtain more detailed information on the effects of increased [K^+^], we recorded intracellularly from pyloric neurons while superfusing 2.5x[K^+^] saline.

### PD and LP neurons depolarize and temporarily lose spiking activity in high extracellular potassium

Intracellular recordings showed a marked loss of spiking activity from pyloric neurons (crash) in response to the application of 2.5x[K^+^] saline, in agreement with the previously described extracellular recordings. In the representative example shown in Figure 2, the PD and LP neurons fired bursts robustly in normal physiological saline (Fig. 2*Ai*). Within a few minutes of the start of 2.5x[K^+^] saline application, the minimum membrane potentials of the PD and LP neurons depolarized by 15 and 22mV respectively (Fig. 2*Aii*), coincident with a reduction in firing frequency. Subsequently, the activity of both the LP and PD neurons became more burst-like over the course of the 90-minute application of 2.5x[K^+^] saline (Fig. 2*Aiii-iv*) and recovered to normal baseline behavior when returned to physiological saline (Fig. 2*Av*).

**Figure 2.**
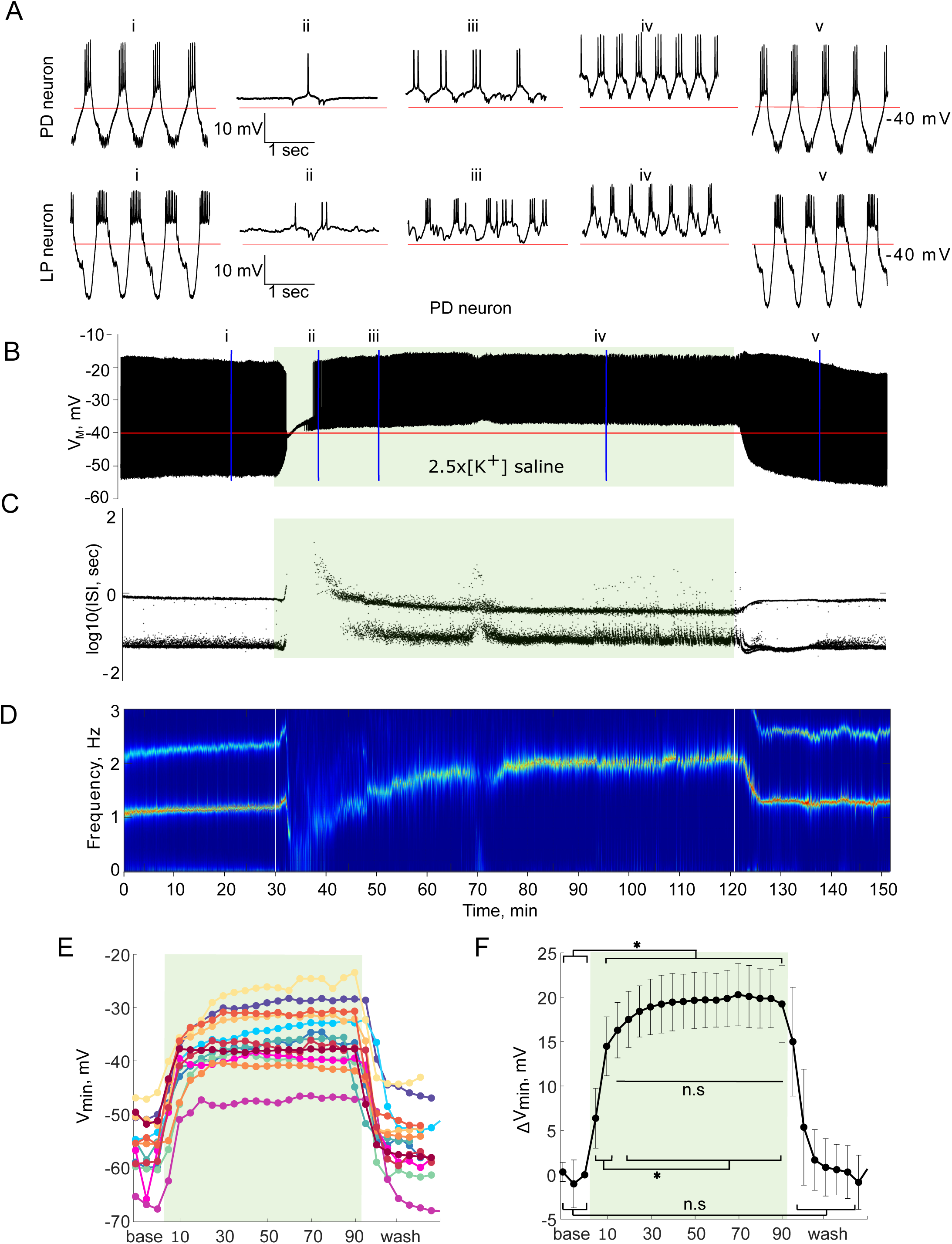
Activity of PD and LP neurons in 2.5x[K^+^] saline. Green shaded boxes indicate the period of 2.5x[K^+^] saline superfusion. (A) Three-second segments of PD and LP activity are shown in physiological saline (i), 10 (ii), 20 (iii) and 70 (iv) minutes into application of 2.5x[K^+^] saline, and upon return to physiological saline (v). (B) Voltage trace of a PD neuron’s activity for the entire representative experiment. (C) Interspike intervals (ISI) of the PD neuron’s activity over the course of 2.5x[K^+^] saline application plotted on a log scale. (D) Spectrogram of the PD neuron’s voltage trace. The color code represents the amplitude density, with red representing the maximum amplitude density and blue the minimum amplitude density. (E) Mean minimum membrane potential for each PD neuron in five-minute bins. (F) Average change in PD neurons’ minimum membrane potential compared to baseline in five-minute bins. Error bars represent standard deviations. Minimum membrane potential depolarized significantly when 2.5x[K^+^] saline was applied (One-way repeated measures ANOVA, Tukey post-hoc, all p < 0.05), but did not change significantly between 10 and 90 minutes in 2.5x[K^+^] (n.s., all p > 0.05). The PD minimum membrane potential returned to baseline levels in wash (n.s., baseline compared to wash, all p > 0.05).

The full pattern of depolarization and recovery of spiking can be seen by plotting time-condensed voltage traces for the PD neuron (the response of the LP neuron closely resembles that of the PD neuron) for the entire 150-minute experiment. In this trace, the membrane potential initially depolarized in 2.5x[K^+^] saline and was followed by a loss of spiking activity (Fig. 2*B*). To visualize spiking behavior over the course of the experiment, we plotted the instantaneous interspike intervals (ISIs) of the PD neuron for the whole experiment on a log scale (log_10_(ISI), Fig. 2*C*). All healthy PD neurons in control saline had regular bursting activity that yielded a bimodal distribution of ISIs, reflecting the relatively longer ISI period between bursts and the shorter ISIs of spikes within a burst (Fig. 2*C*). Over the course of the 2.5x[K^+^] saline application, both the PD and LP neurons recovered rhythmic bursts of action potentials, which is clear from the re-emergence of two ISI bands (Fig. 2*C*).

Bursting activity is suggestive of the re-appearance of slow membrane potential oscillations. These slow oscillations are well visualized in spectrograms of the neuron’s membrane potential. Recovery of bursting activity in elevated [K^+^] saline can be seen as the re-appearance of a strong frequency band at 1-2Hz in the voltage spectrogram (Fig. 2*D*).

We calculated the most hyperpolarized point of the membrane potential in each burst averaged over five-minute bins for all PD neurons to study the depolarizing effect of 2.5x[K^+^] saline over time. PD neurons depolarized upon application of 2.5x[K^+^] saline and remained depolarized for the duration of the application (Fig. 2*E*). Figure 2F shows pooled data from these experiments, PD neurons depolarized upon application of 2.5x[K^+^] saline (Fig. 2*F*, one-way repeated measures ANOVA, F (26,312) = 152.43, Tukey post-hoc, all p<0.05, average depolarization after 10 minutes, 14.5 ± 3.3mV) After the initial change in the first 10 minutes in 2.5x[K^+^], the minimum membrane potential did not change for the remainder of the elevated [K^+^] application (n.s. differences for all comparisons 15 minutes – 90 minutes 2.5x[K^+^]). The minimum membrane potential returned to baseline levels when the preparations were returned to physiological saline (n.s. differences between all baseline and wash timepoints). The behavior of LP neurons in 2.5x[K^+^] was very similar to that of PD neurons; LP neurons depolarized by 16.5 ± 2.9mV after 10 minutes in 2.5x[K^+^] (n = 5) and remained depolarized for the duration of the elevated [K^+^] application.

Although pyloric activity and circuit connectivity are highly conserved across animals, responses of individual PD neurons to 2.5x[K^+^] saline varied substantially across animals. 2.5x[K^+^] saline led to a period of silence in 9 of 13 PD neurons, with a large variability in the duration of silence and extent of recovery across animals (N = 13). The duration of silence of the PD neurons elicited by 2.5x[K^+^] saline application varied from 1 to 62 minutes (10.9 ± 5.8 minutes SD).

We were interested to see whether aspects of baseline activity of each PD neuron influenced the neuron’s time to recovery. Therefore, for each PD neuron, we calculated the mean minimum membrane potential during baseline recordings, the change in membrane potential upon application of 2.5x[K^+^] saline, and the baseline bursting frequency. We found no correlation between any of these values and silence duration for a given PD neuron in 2.5x[K^+^] saline (R^2^=0.203, R^2^=0.104, R^2^=0.012 respectively).

### PD neurons in the isolated pacemaker kernel continue spiking in 2.5x[K^+^] saline

From the previous experiments, it was unclear to what extent the crash and recovery of activity in 2.5x[K^+^] saline was impacted by presynaptic inputs to the PD neurons. The only feedback from the rest of the pyloric circuit to the pacemaker ensemble is the glutamatergic inhibitory input from the LP neuron (Eisen and Marder, 1982). Thus, the pacemaker kernel can be studied in isolation from the other pyloric neurons by superfusing saline with 10^-5^M picrotoxin (PTX), which blocks ionotropic glutamatergic synapses in the STG (Fig. 3*A*)(Bidaut, 1980).

**Figure 3.**
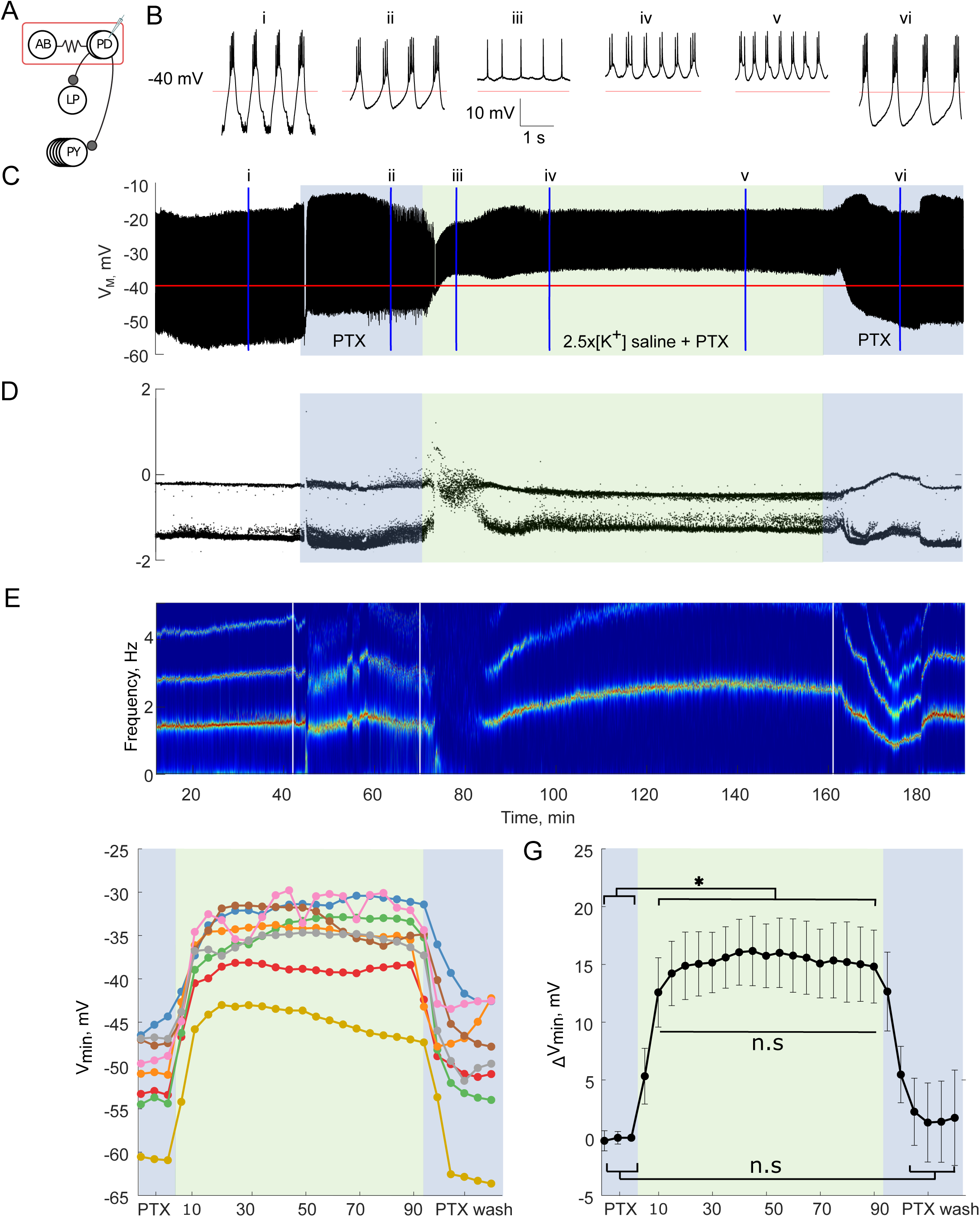
Activity of a PD neuron isolated from its presynaptic glutamatergic synapses with picrotoxin (PTX) in 2.5x[K^+^] saline. (A) Connectivity diagram of the pyloric network with picrotoxin (PTX) blocking inhibitory glutamatergic synapses, and only cholinergic synapses still present. (B) Three-second segments of the PD neuron’s activity in physiological saline (i), 20 min into application of 10^-5^M PTX saline (ii), at 10 (iii), 20 (iv) and 70 (v) minutes into application of 2.5x[K^+^] PTX saline, and upon return to physiological saline (vi). (C) Voltage trace of the PD neuron over the entire experiment. The blue shaded boxes indicate time of 10^-5^M PTX saline superfusion, and the green shaded box indicates the 90-minute period of 2.5x[K^+^] PTX saline superfusion. Color scheme is maintained in D, F, and G. (D) ISIs from the PD neuron over the course of the experiment plotted on a log scale. (E) Spectrogram of PD neuron’s voltage trace. The color code reflects the amplitude density. (F) Mean minimum membrane potential of each PTX PD neuron in five-minute bins. (G) Average change in PTX PD neurons’ minimum membrane potential compared to baseline in five-minute bins. Error bars represent standard deviations. Minimum membrane potential depolarized significantly when 2.5x[K^+^] PTX saline was applied (One-way repeated measures ANOVA, Tukey post-hoc, all p < 0.05) but does not change significantly after 10 minutes in 2.5x[K^+^] PTX for the duration of the elevated [K^+^] application (n.s., all p > 0.05). The minimum membrane potential returned to baseline levels in wash (n.s., baseline compared to wash, all p > 0.05).

The response of PD neurons to 2.5x[K^+^] saline in the presence of PTX was markedly different from the behavior of PD neurons in the absence of PTX. Figure 3 illustrates a representative example of the activity of a PD neuron in 2.5x[K^+^] PTX saline. The neuron initially switched from rhythmic bursting to tonic spiking activity in 2.5x[K^+^] PTX saline (Fig. 3*Biii*), followed by recovery of bursting activity that became more pronounced with time (Fig. 3*Biv – vi*). There was no interruption of spiking activity in PD neurons upon the superfusion of 2.5x[K^+^] PTX saline (Fig. 3*C*) as compared to 2.5x[K^+^] alone (compare to Fig. 2*B*). The recovery of bursting activity in 2.5x[K^+^] PTX saline can also be seen in the emergence of two distinct ISI bands (Fig. 3*D*) and the emergence of a clear frequency band in the spectrogram of the PD intracellular voltage trace (Fig. 3*E*).

We further quantified the responses of PD neurons to 2.5x[K^+^] PTX saline by calculating the mean minimum membrane potential in five-minute bins for each neuron across the experiment (N = 8, Fig. 3*F*). Similar to PD neurons in the intact circuitry, all PD neurons in PTX depolarized in 2.5x[K^+^] saline (Fig. 3*G*, one-way repeated measures ANOVA, F(26,182) = 82.40, Tukey post-hoc, all p < 0.05, mean depolarization after 10 minutes 12.6 ± 3.0mV). The minimum membrane potential of PD neurons in PTX then remained stable until the end of the application of 2.5x[K^+^] and hyperpolarized back to baseline levels when returned to physiological [K^+^] saline (Fig. 3*G*, n.s. differences between all baseline and wash timepoints).

### Synaptic inputs alter response of PD neurons to 2.5x[K^+^] saline

The activity of PD neurons upon initial 2.5x[K^+^] saline application differed in the presence or absence of PTX, which indicates existence of a circuit-driven response to elevated [K^+^] saline. To quantify this effect, we used values from the Hartigan’s dip test calculated from 2-minute bin log(ISI) distributions to determine the time that each PD neuron was either bursting, tonically firing, sparsely spiking, or silent throughout the experiment and plotted the assigned category for each PD neuron over time (see methods).

In the intact circuit, the majority (N = 9 of 13) of PD neurons exhibited a period of silence of at least 2 minutes following the application of 2.5x[K^+^] saline, then recovered spiking activity over a variable amount of time (Fig. 4*A*). In 2.5x[K^+^] PTX saline, PD neurons either remained active or only briefly went silent, then demonstrated robust recovery of spiking or bursting activity (Fig. 4*B*). The average time of PD silence in 2.5x[K^+^] saline was significantly less with the addition of PTX (Fig. 4*C*, Wilcoxon rank-sum test p = 0.013), as was the average time of PD sparse firing between PTX and control PD neurons (Fig. 4*D*, Wilcoxon rank-sum test, p = 0.034). In both the duration of silence and sparse firing states, there is one PD neuron that spent substantially more time in either of these states; however, removing this point did not affect the statistical differences between the control and PTX groups.

**Figure 4.**
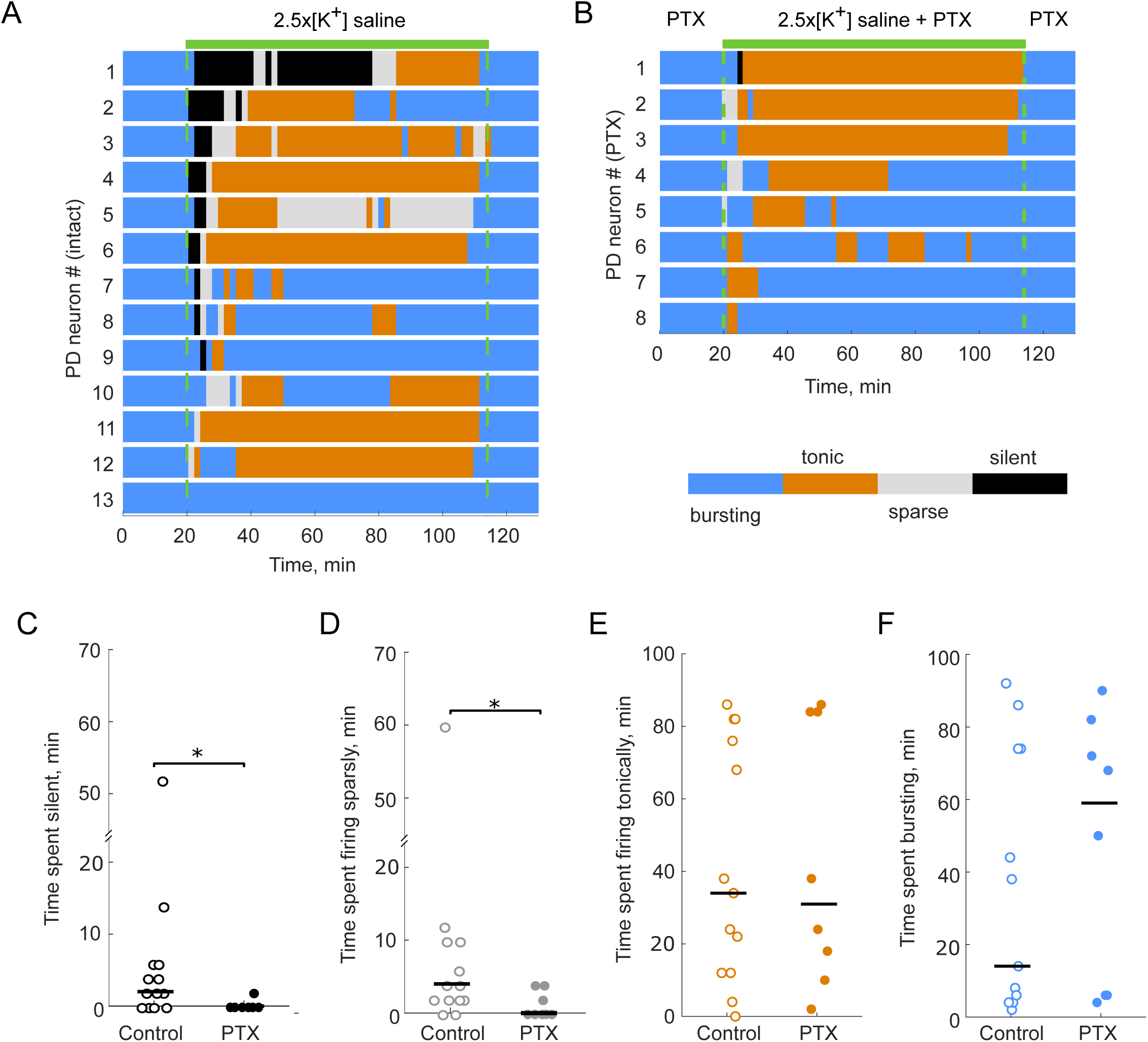
PD neurons respond differently to 2.5x[K^+^] saline in the presence and absence of PTX. (A) Activity patterns (defined in methods and denoted by color blocks) of control PD neurons exposed to 2.5x[K^+^] saline. Each line represents the activity of a single PD neuron. (B) Activity patterns of PTX PD neurons exposed to 2.5x[K^+^] PTX saline. (C) Control PD neurons exhibit longer periods of silence (black) upon 2.5x[K^+^] saline application compared to PD neurons in PTX (Wilcoxon rank-sum test p = 0.0129). (D) Control PD neurons exhibit longer periods of sparse firing (grey) upon 2.5x[K^+^] saline application compared to PD neurons in PTX (Wilcoxon rank-sum test p = 0.034). (E) Control PD neurons do not show significant differences in the amount of time in tonic firing (orange, n.s., p = 0.77) or in (F) burst firing (blue) modes upon 2.5x[K^+^] saline application compared to PD neurons in PTX (n.s., p = 0.51).

Neither the differences in average time of tonic firing (Fig. 4*E*, Wilcoxon rank-sum test, p = 0.77), nor the differences in average time of bursting activity were statistically significant in the presence or absence of PTX (Fig. 4*F*, Wilcoxon rank-sum test, p = 0.51) in 2.5x[K^+^] saline.

### Graded synaptic currents between the LP and PD neurons do not change substantially over time in 2.5x[K^+^]

STG neurons release neurotransmitter as a graded function of presynaptic membrane potential (Graubard, 1978; Manor et al., 1997), with a release threshold close to the trough of the slow wave seen during normal activity. PD neurons temporarily lose and then regain spiking activity in 2.5x[K^+^]. One possible explanation for these results is that graded presynaptic inhibition from the LP neuron, which also temporarily lose activity, onto the PD neuron decreases over time in 2.5x[K^+^] saline, allowing spiking activity to re-appear. We were interested specifically in the role of graded inhibition, because the recovery of spiking activity in PD neurons typically occurs while the rest of the pyloric network is silent.

To test this, we measured the strength of the LP to PD and PD to LP synapses over time in TTX control saline and in 2.5x[K^+^] TTX saline. We measured the synaptic amplitudes in two ways, first using two-electrode current clamp (N = 3) and then using two-electrode voltage clamp (N=3). Both techniques consistently showed that there is no decrease in graded synaptic transmission during adaptation to high extracellular [K^+^]. Figure 5 summarizes the results of the voltage-clamp experiments. In 2.5x[K^+^] TTX saline there was a large decrease in the graded synaptic current from LP to PD measured at -10mV (Fig. 5*A*). This decrease in synaptic current can be accounted for by the increase in extracellular [K^+^] leading to a depolarization of the synaptic reversal potential. The magnitude of the synaptic current in 2.5x[K^+^] saline did not weaken over time, and in the example shown, slightly increased (Fig. 5*B*), indicating that the recovery is not likely a result of transmitter depletion. In all 6 experiments, there was no decrease in the LP to PD synapse strength over time in 2.5x[K^+^] saline (Fig. 5*C* *(voltage clamp)*).

**Figure 5.**
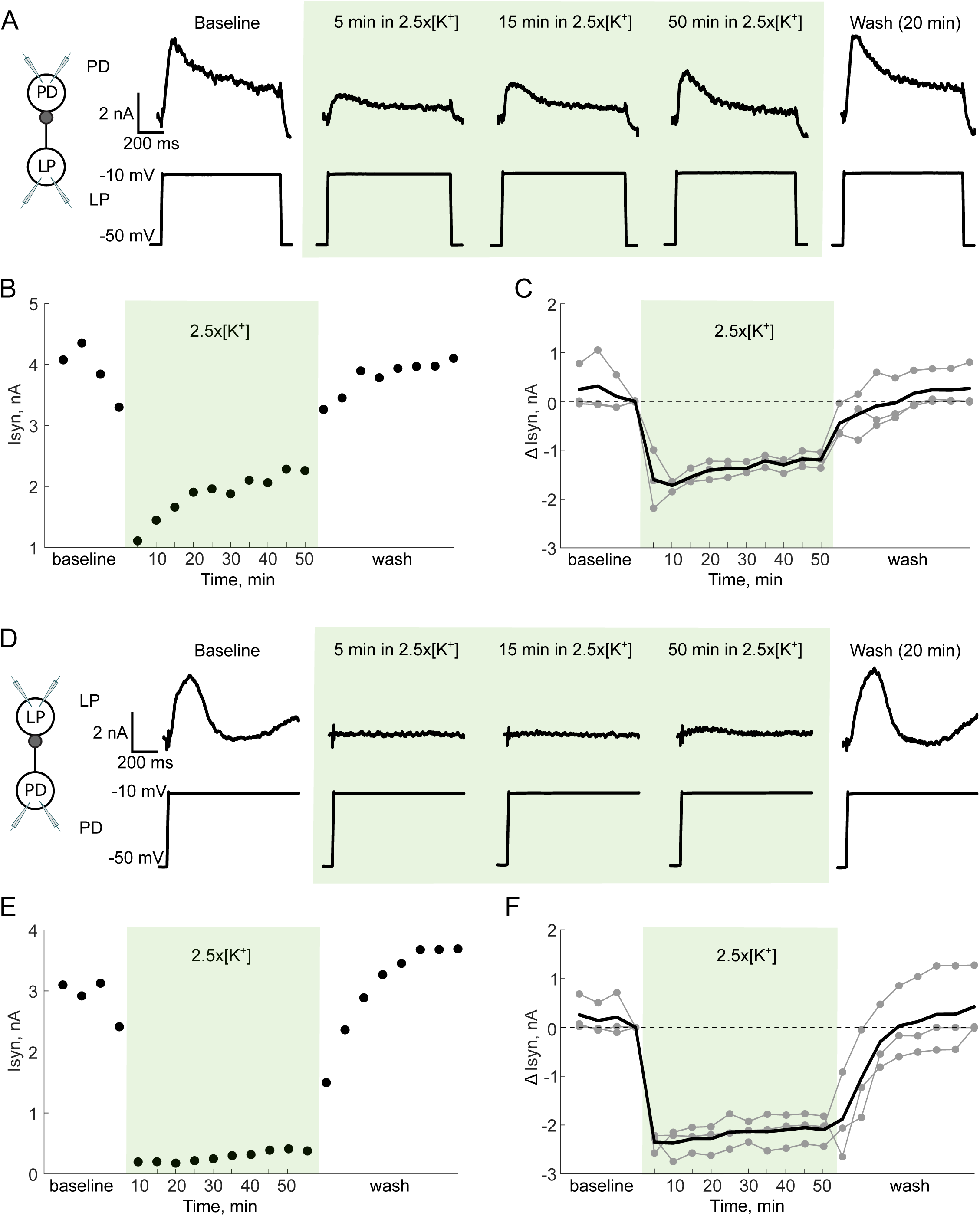
Graded synaptic currents between LP to PD neurons do not change substantially over time in 2.5x[K^+^] saline. Two-electrode voltage clamp in both the LP and PD neurons. The PD neuron’s membrane potential was clamped to either -90mV or -10mV for 3 seconds while stepping the LP neuron from -50mV to - 10mV for 1 second. A-C show the synaptic currents from the LP neuron to the PD neuron, with the PD neuron at -10mV and the LP neuron depolarized to -10mV (A) Representative traces of the synaptic current in a PD neuron in response to the LP neuron’s depolarization over time in 2.5x[K^+^] and wash. (B) Maximum synaptic current elicited in the PD neuron over the course of the entire experiment. (C) Change in synaptic current relative to the last point measured in baseline. Individual experiments are shown in grey, the mean change in current is represented by the black line. D-F show the synaptic currents from the PD neuron to the LP neuron, with the LP neuron was held at -50mV and PD neuron depolarized to -10mV (D) Representative traces of the synaptic current in the LP neuron in response to the PD neuron’s depolarization over time in 2.5x[K^+^] and wash. (E) Maximum synaptic current elicited in the LP neuron over the course of the entire experiment. (F) Change in synaptic current elicited in the LP neuron relative to the last point measured in baseline. Individual experiments are shown in grey, the mean change in current is represented by the black line.

In voltage clamp, we also measured the strength of graded synaptic transmission from the PD neuron to LP neuron (Fig. 5*D*). The magnitude of the synaptic current also decreased markedly upon application of 2.5x[K^+^] saline (again because the reversal potential depolarized) and remained low throughout the application (Fig 5*E*). This effect was consistent across all preparations (Fig. 5*F*). Thus, decreased graded inhibition cannot account for the recovery of spiking activity in pyloric neurons over time in elevated [K^+^] saline.

### Intrinsic excitability of PD neurons changes rapidly during exposure to 2.5x[K^+^] saline

To confirm that the periods of silence in high K^+^ were not due to depolarization block, we applied 60-second steady ramps of current from -4nA to +2nA to PD neurons at several time points during the period of silence elicited by application of 2.5x[K^+^] saline. In silent PD neurons, action potentials could always be induced by injecting positive current, indicating that this period of silence is not due to depolarization block (Fig. 6*A*, ramps at 5-minutes and 10-minutes).

**Figure 6.**
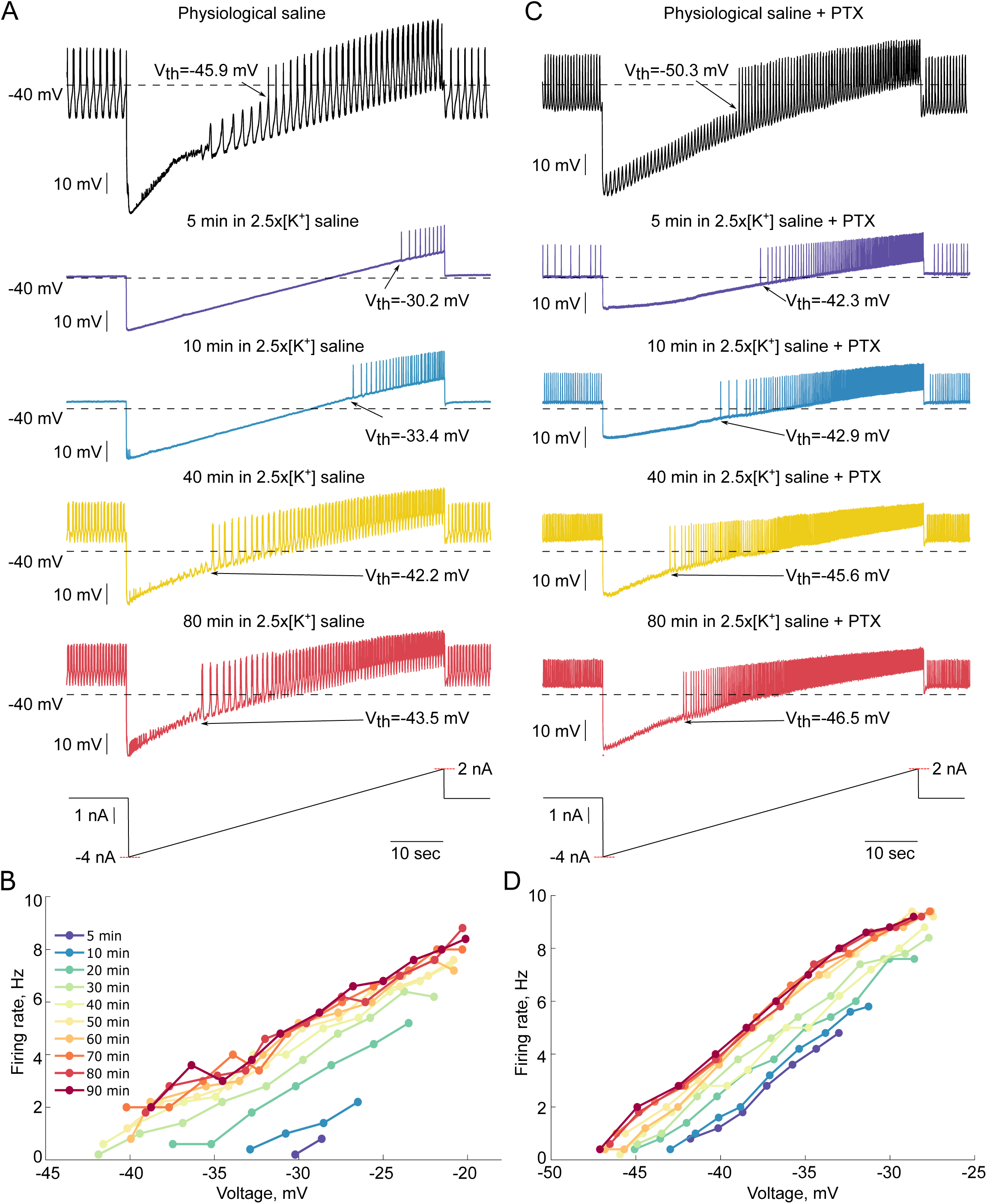
Excitability of representative PD neurons during exposure to 2.5x[K^+^] saline. Two-electrode current clamp was used to inject current ramps from - 4nA to +2nA over 60 seconds. (A) Representative activity during ramps from a PD neuron in 2.5x[K^+^] at 5, 10, 40, and 80 minutes after the onset of 2.5x[K^+^] saline application. (B) Average firing rates calculated in 5-second bins for each ramp in 2.5x[K^+^] saline from the neuron shown in A plotted against the average membrane potential of the corresponding bin. (C) Representative activity during ramps for a PD neuron in PTX saline at 5, 10, 40 and 80 minutes after the onset of 2.5x[K+] saline application. (D) Average firing rates calculated in 5-second bins for each ramp in 2.5x[K^+^] saline from the neuron shown in C are plotted against the average membrane potential in the corresponding bin.

To determine the excitability of PD neurons during application of 2.5x[K^+^], we repeated the slow current ramps from -4nA to +2nA at 5, 10, 20, 30, 40, 50, 60, 70, 80, and 90 minutes after the beginning of the 2.5x[K^+^] saline application in the presence or absence of PTX. In the representative example shown in Figure 6*A*, there was a clear change in the spike threshold and frequency of spikes elicited in the PD neuron by the current ramp as a function of time in 2.5x[K^+^] saline. As more time in 2.5x[K^+^] elapsed, more spikes were elicited at the same membrane potentials during the current ramp (Fig. 6*B*).

Similar to PD neurons in the intact circuitry, PD neurons in the presence of PTX also a more excitable over time in 2.5x[K^+^] saline. In the representative example shown in Figure 6*C*, although the PD neuron remained tonically active throughout the application of 2.5x[K^+^] saline, there was a shift in both the spike threshold and spike frequency during the ramps over time in 2.5x[K^+^]. Similar to the PD neuron in the intact circuitry (Fig. 6*B*), the firing rate of the PD neuron in PTX increased over time in 2.5x[K^+^] at all membrane potentials during the ramps (Fig. 6*D*).

Figure 7 summarizes the changes in spike threshold and firing rate of PD neurons across all preparations as a function of time in 2.5x[K^+^] saline. In both control and PTX conditions, PD neuron spike threshold hyperpolarized over time in 2.5x[K^+^] saline (Fig. 7*A*). For most of these neurons the greatest changes in firing rate and spike threshold occurred during the first 10 minutes of the 2.5x[K^+^] saline application, which is similar to the time in which most PD neurons recovered spiking activity in elevated [K^+^]. Control PD neurons had significantly different spike thresholds between the 5-minute and the 30 through 90-minute time points in 2.5x[K^+^] saline and also between the 10-minute and the 70 through 90-minute time points in 2.5x[K^+^] saline (Fig. 7*A*, one-way repeated measures ANOVA, Tukey Post-Hoc test, F(14,56) = 19.99, all p < 0.05). Across all PTX PD neurons there was a significant change in spike threshold between the 5-minute and 20 through 90 minute time points in 2.5x[K^+^] saline (Fig. *7A*, one-way repeated measures ANOVA, Tukey Post-Hoc test, F(14,98) = 30.98, all p > 0.05).

**Figure 7.**
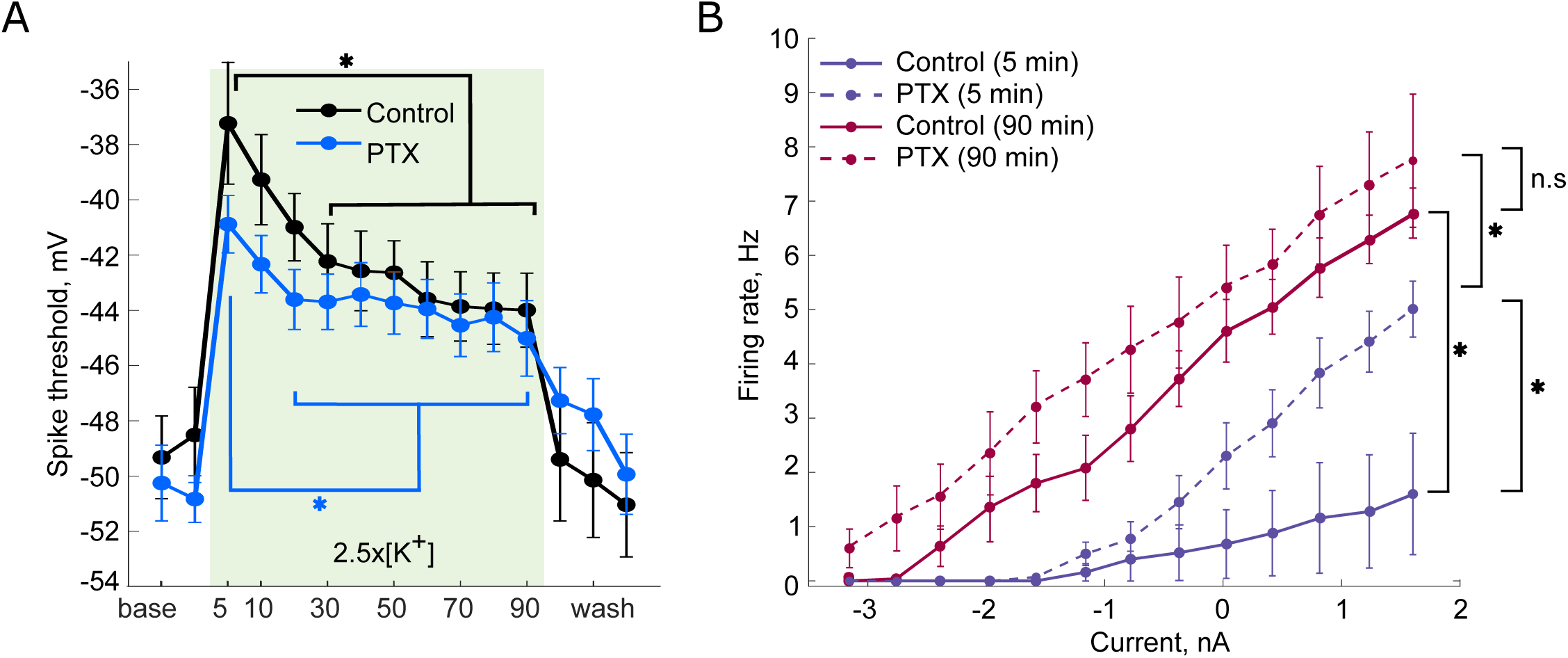
Intrinsic excitability of PD neurons changes in 2.5x[K^+^] saline. (A) Spike thresholds (mean+-sem) for control and PTX PD neurons, calculated from 60-second ramps from -4nA to +2nA. One-way repeated measures ANOVA with Tukey Post-Hoc test (* = p < 0.05, all comparisons between brackets). (B) Average F-I curves for control PD neurons (solid lines) and PD neurons in PTX (dashed lines) in the beginning of 2.5x[K^+^] saline application(5-minute, blue lines) and at the end of 2.5x[K^+^] saline application (90-minute, red lines). There was a significant difference in the F-I curves of PTX and control PD neurons after 5 minutes in 2.5x[K^+^] saline (Two-way repeated measures ANOVA, p = 0.001). Both control and PTX PD neurons become more excitable between the early (5-minute) and late (90-minute) time points in 2.5x[K^+^] (Two-way repeated measures ANOVA, p = 0.009 and p = 0.006 respectively).

To visualize changes in PD neuron excitability and how this depends on both synaptic inputs and time in 2.5x[K^+^] saline, we calculated average Frequency-current (F-I) curves for all PD neurons at the beginning (5-minute ramp) and at the end (90-minute ramp) of the 2.5x[K^+^] saline application. Early in the application (5 minutes) of 2.5x[K+], PTX PD neurons showed higher firing rates than control PD neurons (Fig 7*B*, purple lines, two-way repeated measures ANOVA, F(1,11) = 18.52, p = 0.001). This difference represents the initial effect of local inhibitory connections on neuronal excitability and is consistent with the fact that in the presence of PTX, neurons remained active upon application of 2.5x[K^+^], whereas control PD neurons typically lost spiking activity. There was a shift in the average F-I curve between the early (5-minute) and late (90-minute) time points in 2.5x[K^+^] saline (Fig. 7*B* solid lines, two-way repeated measures ANOVA, F(1,4) = 23.04, p = 0.009) indicating an increase in excitability. This was also seen for PTX PD neurons in 2.5x[K^+^]. There was a statistically significant difference between the early (5-minute) and late (90-minute) PTX PD neuron F-I curves (Fig 7*B* dashed lines, two-way repeated measures ANOVA, F(1,7) = 15.22, p = 0.006). Finally, by the end of the 2.5x[K^+^] application (90-minutes), there was no significant difference between the F-I curves of PD neurons in the presence or absence of PTX (Fig 7*B* red lines, two-way repeated measures ANOVA, F(1,11) = 1.42, p = 0.258).

## Discussion

External solutions containing high extracellular [K^+^] are commonly used to depolarize neurons and other tissues, often with the assumption that depolarization will be associated with an increase in neuronal activity. Some treatments with high extracellular [K^+^] are done transiently, for short periods of time, whereas in other experiments, preparations are kept in high extracellular [K^+^] for many hours. In this paper we demonstrate adaptations to high [K^+^] that occur in minutes to hours, that could complicate the interpretation of some experiments that use elevated [K^+^]. Moreover, the phenomena described here may also have implications for disease states in which high extracellular K^+^ concentrations are abruptly elevated for minutes, hours or days.

Figure 8 summarizes the core data and conclusions of this paper and illustrates the evolution of the changes in normal and high [K^+^]. Most importantly, this schematic shows that the effects of high [K^+^] are not stationary, but instead have an initial set of effects followed by an adaptation phase. This result suggests that researchers using other preparations should be alert to the possibilities of different short and long-term effects of changing extracellular K^+^ concentrations.

**Figure 8.**
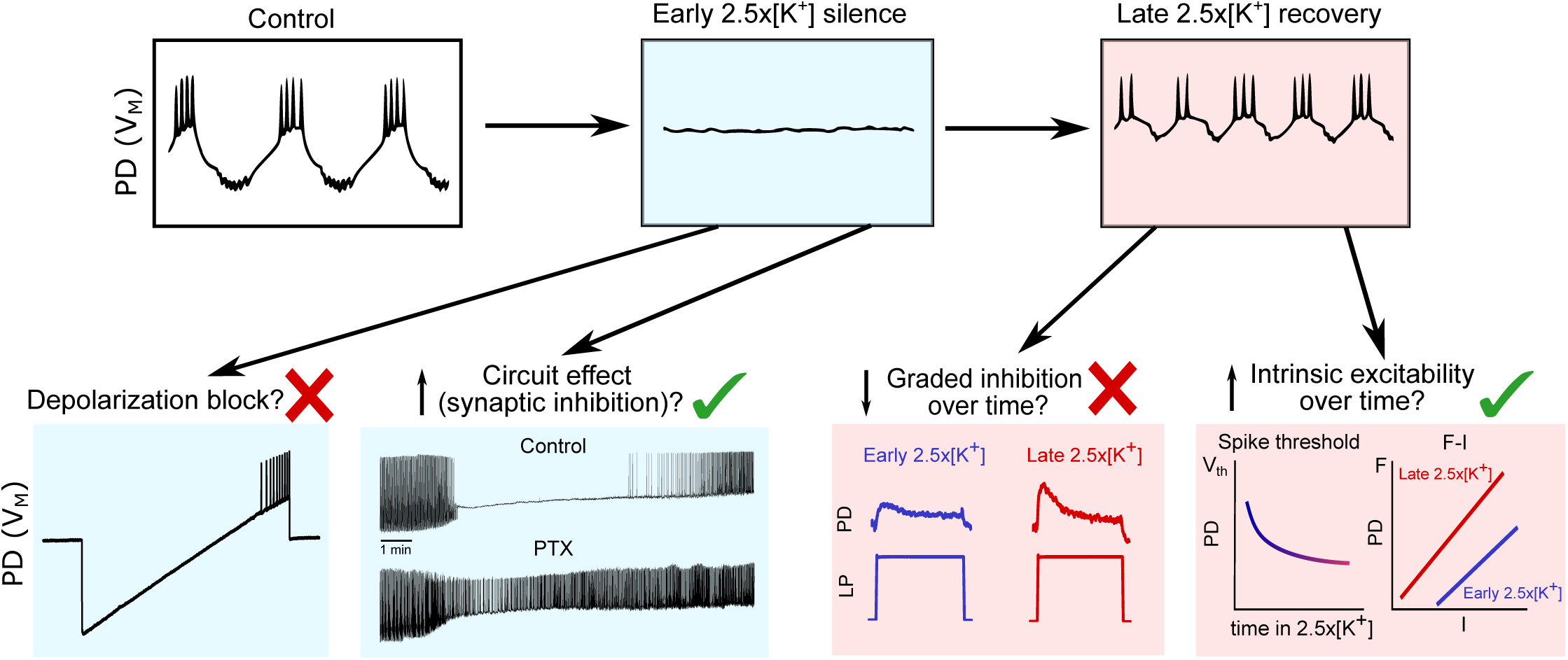
Schematic of possible mechanisms explaining the response to elevated [K^+^]. Boxes on the top row illustrate typical behavior of a PD neuron in control saline, early 2.5x[K^+^] saline (blue) and late 2.5x[K^+^] (pink). The initial loss of spiking activity in early 2.5x[K^+^] saline could potentially be explained by depolarization block, or by circuit mediated inhibition. Action potentials can be elicited in silent PD neurons by injecting positive current, showing that this silence is not due to depolarization block (Red X, trace from Fig. 6). PD neurons in early 2.5x[K^+^] saline do not lose spiking activity when inhibitory glutamatergic synapses are blocked, indicating that the initial silence can be explained by local inhibition from the LP neuron (Green ✔, traces from Fig. 2 and 3). The second phenomenon is the recovery of spiking and bursting activity late in 2.5x[K^+^] saline application. This recovery of spiking could be due to a reduction in graded inhibition over time and/or an increase in intrinsic excitability. Upon application of 2.5x[K^+^] saline, the response of PD neurons to LP depolarization is greatly reduced, and slightly increases as time increases in 2.5x[K^+^] saline indicating that a change in synaptic inhibition is unlikely to be responsible for the recovery of spiking activity (Red X, traces from Fig. 5). Measurements of spike threshold and F-I curves from PD neurons late and early in 2.5x[K^+^] saline show that the intrinsic excitability of PD neurons increases over time (Green ✔, data from Fig. 7). This shift in excitability could be responsible for the recovery of spiking and rhythmic activity over time.

### Inhibitory effect of a depolarizing stimulus

It is generally assumed that positive current or depolarization of a neuron’s membrane potential will lead to an increase in neuronal activity and that extreme depolarizations can lead to loss of activity through depolarization block. However, we present a case in which depolarization due to elevated extracellular [K^+^] instead leads to a transient network silence that is not due to depolarization block (Fig. 8, blue panels).

In the pyloric circuit, all local synaptic connections are inhibitory (Eisen and Marder, 1982; Miller and Selverston, 1982; Rosenbaum and Marder, 2018). Synaptic transmission in the STG is both graded and spike-mediated, meaning that action potentials are not required for inhibitory synapses to function (Graubard et al., 1980; Manor et al., 1997). Therefore, the observed decrease in both bursting and spiking activity in elevated extracellular [K^+^] could be caused by global depolarization leading to increased inhibition that initially suppresses spiking and bursting activity. In support of this theory, elevated [K^+^] did not have the same inhibitory effect on PD neurons with glutamatergic inhibitory synapses blocked compared to PD neurons with intact synaptic connections. We speculate that similar blockades of circuit activity could occur whenever synaptic inhibition is strengthened. Moreover, in studies of proprioceptive neurons of the blue crab and the muscle receptor organ of the crayfish, increased [K^+^] also has an inhibitory effect at concentrations not thought to cause a depolarization block (Malloy et al., 2017).

### Adaptation to global perturbation

The PD/AB pacemaker unit of the pyloric circuit exhibited rapid adaptation to the disruptive stimulus of increased extracellular [K^+^] (Fig. 8, pink panels). Although the triphasic pyloric rhythm was not fully restored in 2.5x[K^+^] saline, PD neurons exhibited rapid changes in excitability over several minutes, which corresponded to the recovery of spiking and, in many cases, bursting activity. This time course is similar to a fascinating series of homeostatic mechanisms in *Drosophila* that also occurs over several minutes (Frank et al., 2006).

The time course of adaptation in the present study contrasts with many examples of homeostatic plasticity in which loss of activity elicits changes in gene expression to restore activity that are thought to occur over hours to days (Desai et al., 1999; Cudmore and Turrigiano, 2004; Turrigiano, 2012). Such mechanisms can be induced by changes in extracellular [K^+^]. Rat myenteric neurons cultured in elevated [K^+^] serum for several days exhibit long-lasting changes in Ca^2+^ channel function (Franklin, 1992). Similarly, culturing rat hippocampal pyramidal cells in high [K^+^] medium for several days leads to activation of calcium-dependent changes in the intrinsic excitability of the neurons that can be seen in changes in the input resistance and the resting membrane potential (O’Leary et al., 2010).

Changes in K^+^ channel densities are also associated with homeostatic regulation of neuronal activity. Depolarization of crustacean motor neurons with current pulses for several hours alters K^+^ channel densities in a cell-specific manner through a calcium-dependent mechanism (Golowasch et al., 1999). In addition, when inhibitory GABA receptors are blocked for several days in cerebellar granule cells, they maintain their responses to excitatory input by strengthening voltage-independent K^+^ conductances (Brickley et al., 2001). Adaptation to global perturbation over several hours to days is well described by computational models via calcium signals that influence expression levels of ion channels (O’Leary et al., 2014; O’Leary, 2018).

In the STNS, the upstream commissural ganglia and the esophageal ganglion release a wide range of neuromodulators onto the STG that affect excitability of cells and the pyloric rhythm (Marder and Bucher, 2007; Marder, 2012). Pyloric neurons in the STG also exhibit a form of long-term adaptation in response to removal of these modulatory inputs. Some preparations in which neuromodulatory inputs are removed initially lose rhythmicity and gradually recover over the course of several days (Thoby-Brisson and Simmers, 1998, 2002; Luther et al., 2003; Gray and Golowasch, 2016; Gray et al., 2017). Nonetheless, when preparations remain active after removal of neuromodulatory inputs, they tend to maintain a level of activity similar to that shown in preparations that recover after losing activity (Hamood et al., 2015), suggesting that there may be a target activity level for the circuit.

The rapid adaptation of PD neurons in elevated extracellular [K^+^] is most likely due to changes in cell intrinsic conductances (Fig. 8, pink panel). In crustacean motor neurons, cell-intrinsic changes in current densities have been shown to be modulated by second-messenger kinase pathways activated by global depolarization (Ransdell et al., 2012). Rapid changes in circuit state could also be influenced by neuromodulation, as the experiments in this study were conducted on the entire STNS, including connections from the upstream modulatory ganglia that were also exposed to elevated [K^+^].

### Variable responses to similar perturbations

Within the STG, conductance densities and strengths of synaptic connections can vary 2 to 6 fold in magnitude between individuals (Schulz et al., 2006; Schulz et al., 2007; Goaillard et al., 2009; Shruti et al., 2014; Temporal et al., 2014; Alonso and Marder, 2019). Variability in neuronal conductances underlying similar activity patterns has also been demonstrated across phyla (Swensen and Bean, 2005; Nelson and Turrigiano, 2008; Roffman et al., 2012; Tran et al., 2019). In addition, computational modeling of the pyloric network has revealed that multiple combinations of parameters can give rise to similar activity patterns (Goldman et al., 2001; Prinz et al., 2004; Taylor et al., 2009; O’Leary et al., 2014; Marder et al., 2015).

Circuits and individual neurons with apparently identical outputs under control conditions can present distinctly different responses to perturbation due to underlying differences in network parameters (Tang et al., 2012; Hamood and Marder, 2014; Haddad and Marder, 2018; Haley et al., 2018; Alonso and Marder, 2019). In this study we observed a range in the time course of the adaptive response to the same perturbation, which may suggest individual parameter differences that are not evident in control conditions. The application of elevated [K^+^] saline to identified STG neurons provides additional evidence that differences in individual conductance parameters can influence responses to global perturbation (Alonso and Marder, 2019).

### Potential implications for disease states

Hyperkalemia is associated with a number of human disease states. Chronic Kidney Disease (CKD) leads to increases in serum [K^+^] up to three times the normal levels, directly affecting neuronal excitability and changes in neuronal properties (Krishnan and Kiernan, 2009; Arnold et al., 2014). Similarly, increased activity of a group of neurons can increase the extracellular [K^+^] in the surrounding tissue (Baylor and Nicholls, 1969; Kříž et al., 1974) and epileptic seizures and brain trauma can lead to increases in [K^+^] in surrounding brain regions (Moody et al., 1974; Katayama et al., 1990; Silver and Erecinska, 1994; Fröhlich et al., 2008). In this study we showed that changes in [K^+^] similar to those reported in pathological conditions produce not only immediate change in the activity of STG motor neurons, but also long-lasting changes in the intrinsic properties of these neurons. Transient changes in extracellular [K^+^] have been shown to cause long lasting changes in the organization and phosphorylation pattern of K^+^ channels (Misonou et al., 2004), which could lead to long-lasting changes in circuit function (Somjen, 2001, 2002; Rodgers et al., 2007).

### Reassessing global perturbation

The results of this study highlight the lack of a consistent response to a seemingly simple perturbation. In this classical manipulation of increased extracellular [K^+^], we observed a paradoxical decrease in the activity of PD neurons upon bath application of high [K^+^], followed by a recovery of activity in a short period of time. Despite knowing the network connectivity, circuit properties and expected behavior of identified neurons within the STG, we were still unable to *a priori* predict or fully explain the effects of increased [K^+^] on circuit performance. The complex interaction between circuit level effects and cell intrinsic responses to simple changes in ion concentrations underscores the importance of assessing and reporting neuronal activity during such manipulations in any experiment.

## Acknowledgments

Supported by NIH R35 NS097343 and R90 DA033463. The authors thank Daniel Shin for assistance with dissections, and Dr. Stephen Van Hooser for statistical advice. Drs. Sonal Kedia and Jason Pipkin gave us comments on the manuscript.

## Notes

The authors declare no conflicts of interest.

